# Normative models combining fetal and postnatal MRI data to characterize neurodevelopmental trajectories during the transition from in- to ex-utero

**DOI:** 10.1101/2024.03.07.583908

**Authors:** A. Mihailov, A. Pron, J. Lefèvre, C. Deruelle, B. Desnous, F. Bretelle, A. Manchon, M. Milh, F. Rousseau, G. Auzias, N. Girard

**Affiliations:** Institut de Neurosciences de la Timone, UMR 7289, CNRS, Aix-Marseille Université, 13005, Marseille, France; Aix-Marseille Univ, APHM, service de neurologie pédiatrique, Hôpital de la Timone, 13005, Marseille, France; Aix-Marseille Univ, APHM, Service de Gynécologie Obstétrique, Hôpital Nord, 13015, Marseille, France; Aix-Marseille Univ, APHM, Service de Neuroradiologie, Hôpital de la Timone, 13005, Marseille, France; IMT Atlantique, LaTIM U1101 INSERM, 29200, Brest, France

## Abstract

The perinatal period involves transitioning from an intra- to an extrauterine environment, which requires a complex adaptation of the brain. This period is marked with dynamic and multifaceted cortical changes in both structure and function. Most studies to date have focused either on the fetal or postnatal period, independently. To the best of our knowledge, this is the first neurodevelopmental study targeting the cortical trajectory of typically developing perinatal subjects, combining MRIs from both fetal and postnatal participants. Prior to analysis, preprocessing and segmentation parameters were harmonized across all subjects in order to overcome methodological limitations that arise when studying such different populations. We conducted a normative modeling analysis on a sample of 607 subjects, age ranged 24 to 45 weeks post-conception, to observe changes that arise as participants traverse the birth barrier. We observed that the trajectories of global surface area and several volumetric features, including total gray matter, white matter, brainstem, cerebellum and hippocampi, follow distinct but continuous patterns during this transition. We further report three features presenting a discontinuity in their neurodevelopmental trajectories as participants traverse from a fetal to a postnatal environment: the extra-cerebrospinal fluid volume, the ventricular volume and global gyrification. The current study demonstrates the presence of unique neurodevelopmental patterns for several structural features during the perinatal period, and confirms that not all features are affected in the same way as they cross the birth barrier.

**SIGNIFICANCE STATEMENT:** The perinatal phase comprises the fetal and immediate postnatal period, and is generally described as the time surrounding birth. Comprehensively understanding this period is crucial due to the presence of dynamic and multifaceted brain changes. What makes this investigation unique is that it is the first neurodevelopmental study, to the best of our knowledge, focused on the cortical trajectory of typically developing perinatal subjects through the combination of both fetal and postnatal participants into one analysis. We report that certain brain feature trajectories change drastically as fetuses become newborns, while other features remain continuous. These observations are relevant in both the isolation of biomarkers for later cognitive and physiological disorders and in the understanding of typical cerebral development.

## 1. Introduction

The perinatal period is generally defined as the time shortly after conception up until a few weeks to one year after birth (Helfer, 1987; World Health Organization, 2016). Characterizing brain development during this period is challenging as a consequence of numerous complex and intricate genetic, molecular and cellular mechanisms that act in concert during brain maturation (Kostović et al., 2019). This period is particularly important in the isolation of cortical biomarkers since alterations in trajectories and neural patterns of both pre- and postnatal subjects have been associated with later neurodevelopmental and physiological disorders (Girard and Huisman, 2005; Ment et al., 2009; Holland et al., 2014; Bouyssi-Kobar et al., 2016; De Asis-Cruz et al., 2022; Sadhwani et al., 2022; Walhovd et al., 2023). However, most of these studies consider fetal and postnatal periods separately, such that the specific transition from in-utero and ex-utero life remains poorly characterized.

### 1.1. Proper modeling of age effects in early brain development

The widespread commonality of applying linear regression models for estimating the temporal evolution of any feature seems insufficient to understand complex early neurodevelopment. Studying the typically developing cortex during this dynamic developmental phase therefore necessitates the use of proper nonlinear models with larger sample sizes to accurately capture fine-grained spatiotemporal trajectories (Kyriakopoulou et al., 2017; Bethlehem et al., 2022). The normative modeling framework has been proposed as an adequate statistical approach to investigate the nonlinear influence of age on MRI-extracted brain features. It characterizes heterogeneity within a population by considering non-linearity and interactions across several factors such as age and sex (Marquand et al., 2016). Contrary to the case-control paradigm, normative modeling enables the estimation of deviation from a reference sample by focusing on individual-level statistics (Marquand et al., 2019; Gratton et al., 2020). This approach was adopted by a recently developed initiative, *Brain Charts*, that aimed to produce reference charts of cortical morphology evolution across age, starting in the fetal phase up to 100 years old, using over 100 000 subjects (Bethlehem et al., 2022). Despite the remarkable amount of integrated data, the perinatal period is not well represented. More importantly, a closer look at their nonlinear curves indicates a sharp variation in many brain features upon transitioning from a fetal to a postnatal state, which may be interpreted as a discontinuous trajectory. Since such a degree of biological discontinuity in tissue growth is questionable, the origin of these sharp variations is likely related to methodological limitations such as differing segmentation and preprocessing techniques.

### 1.2. Contributions

In the current study, we shed light on normative patterns of perinatal neurodevelopment in several neuroanatomical attributes and provide, to the best of our knowledge, the first empirical test of neurodevelopmental trajectories as they traverse the birth barrier. We characterize patterns of cortical development across several features including global gray matter, white matter, ventricular, extra-cerebrospinal fluid (eCSF), hippocampal, cerebellar and brainstem volumes, as well as surface area and gyrification. We show strong evidence of discontinuity for eCSF, ventricles and gyrification, while observing continuous patterns in the remaining features. We fill a gap within the literature by combining fetal and at-term postnatal subjects into one study, versus investigating them separately as has been generally done until now. By running neurodevelopmental modeling on combined fetal to postnatal subjects, we not only quantitatively characterize important perinatal patterns, but likewise address methodological limitations by applying unified segmentation and preprocessing techniques across all subjects to manage inherent MRI biases. This prevents biases that skew data and can affect results interpretation, and/or introduce discontinuities in the data as seen in the transition between fetal and postnatal populations in the *Brain Charts* paper (Bethlehem et al., 2022).

## 2. Materials and methods

In this section, we outline the criteria for selecting fetal and postnatal participants and the pipelines used for processing this perinatal MRI data. We emphasize the unified segmentation method used for segmenting both fetal and postnatal MRI data and for generating an inner cortical surface interface between cortical gray and white matter. Extensive quality assessments of each step are also described.

### 2.1. The fetal dataset: MarsFet

We constituted our fetal dataset by retrospective access to MRI data acquired during routine clinical appointments at la Timone Hospital in Marseille between 2008 and 2021. Over this period, more than 800 cerebral MRI sessions, spanning a developmental period of 20 to 37 weeks post-conception (wPC), were administered following an indication requested by the Multidisciplinary Center for Prenatal Diagnosis, in the context of usual obstetric assessment during pregnancy. This study was approved by the local ethical committee from Aix-Marseille University (N°2022-04-14-003). We focused on the cerebral MRI sessions acquired using two particular MRI Siemens scanners (Skyra 3T and SymphonyTim 1.5T), using a T2 weighted (T2w) half Fourier single shot turbo spin echo (HASTE) sequence. The details of the MRI acquisition settings are provided in Table 1. To define our fetal normative cohort, close collaboration between contributing medical doctors enabled the design of a set of exclusion criteria combining neuroradiology, obstetrics and pediatric neurology aspects (Table 2).

**Table 1:** A detailed table summarizing MRI acquisition settings for the fetal dataset, MarsFet.

|  | Fetal Dataset |  |  |  |  |  |  |
| --- | --- | --- | --- | --- | --- | --- | --- |
| Scanner Model | Skyra |  |  |  | SymphonyTim |  |  |
| Magnetic Field Strength [Tesla] | 3.0 |  |  |  | 1.5 |  |  |
| Resolution [mm<sup>3</sup>] | 0.6*0.6*3.0 | 0.7*0.7*3.0 | 0.8*0.8*3.0 | 0.7*0.7*3.6 | 0.7*0.7*3.0 | 0.7*0.7*3.4 | 0.7*0.7*3.5 |
| Repetition Time (Tr)[ms] | 4060<br>+- (609) | 3418<br>+- (922) | 3315<br>+- (911) | 3550<br>+- (71) | 1740 | 1690 | 1689<br>+- (59) |
| Echo Time (Te)[ms] | 180<br>+- (1) | 177<br>+- (0) | 177<br>+- (1) | 177<br>+- (1) | 141 | 138 | 131<br>+- (1) |

**Table 2:**
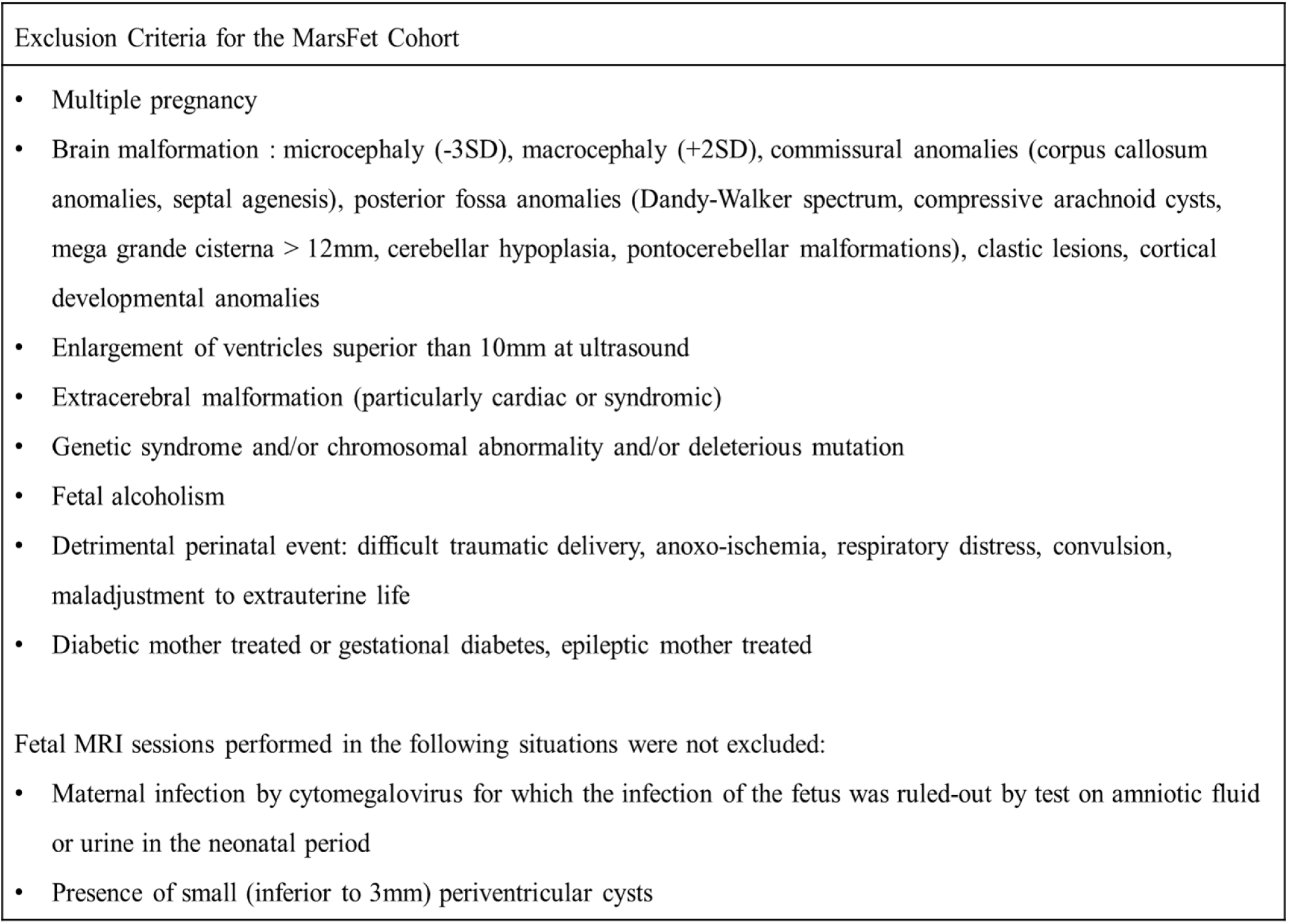
Detailed inclusion/exclusion criteria.

Fetal MRI acquisitions consisted in several 2D T2w images with varying acquisition directions (such as coronal, sagittal, axial, or transverse), referred to as *stacks.* Fetal MRI stacks were denoised using the non-local means approach (Manjón et al., 2010) implemented in the ANTS software (Avants et al., 2009). A mask of the brain was computed for each 2D image using Monaifb (Ranzini et al., 2021). The denoised MRI stacks were corrected for three-dimensional fetal brain intensity bias using the N4 method (Tustison et al., 2010) implemented in ANTS. A 3D high resolution T2w MRI volume (0.5mm iso) was then estimated from the pre-processed MRI stacks using NESVOR v0.2 (Xu et al., 2022) (https://github.com/daviddmc/NeSVoR). The quality of the resulting 3D high resolution T2w volume can be affected by various inaccuracies or artifacts related to the quality of the initial stacks. Since the 3D volume quality in turn impacts the accuracy of the segmentation, we visually inspected each reconstructed volume as detailed in section 2.3.5 below and excluded data of insufficient quality.

### 2.2. The postnatal dataset - developing Human Connectome Project

We used the third release of the publicly available developing Human Connectome Project (dHCP) neonatal dataset (https://www.developingconnectome.org) that consists of 887 MRI scans from 783 infants, spanning a developmental period of 26 to 45 weeks post-conception (Edwards et al., 2022). This dataset thus contains the data from 200 babies born preterm (before 37 wPC). As detailed in Edwards et al. (2022), T2 weighted (T2w) multi-slice fast spin echo (FSE) scans were acquired using a 3T Philips Achieva scanner with in-plane resolution 0.8 × 0.8 mm2 and 1.6 mm slices. We relied on the 3D isotropic (0.5 mm iso) MRI scans reconstructed by combining multi-slice scans after slice-to-volume motion correction using the Cordero-Grande et al. (2016) method. From this dataset, we excluded the MRI scans exhibiting clinical anomalies or presenting incidental findings that were likely to affect further processing (dHCP radiological score of 3, 4 or 5). Participants of a multiple pregnancy were also excluded from our postnatal normative cohort. We further detail the refinement of our final sample in section 2.3.5.

### 2.3. Unified MRI processing pipeline and quality assessment

A unified MRI segmentation and surface extraction pipeline is crucial for quantitatively depicting the development of brain structures as they transition from an in-utero and an ex-utero environment. One key contribution of this work is showcasing the practicability of processing cerebral MRI scans from fetuses and neonates using identical tools, as illustrated in Figure 1. To achieve this, our approach entails combining the state-of-the-art nn-Unet framework (Isensee et al., 2020) with a large amount of T2w postnatal MRI data sourced from the dHCP dataset, in order to transfer a segmentation model from postnatal data to fetal imaging acquired in clinical settings. This approach has two key advantages: 1- the segmentation model is trained on top-quality data, that is not accessible to antenatal acquisitions; 2- the anatomical nomenclature and segmentation accuracy is unified, meaning that defined variations of segmented structures between fetal and postnatal datasets are minimized, contrary to Bethlehem et al. (2022) where different segmentation models were used. Our approach included the following steps: 1) train a state-of-the-art deep-learning 3D segmentation model on the youngest infants from the postnatal dataset (dHCP) from which high quality images and segmentations were available, resulting in the *Unet-preterm* segmentation model; 2) Fine-tune the Unet-preterm model on a fetal ground truth, resulting in the Unet-fetus segmentation model; 3) Segment the rest of the postnatal MRI scans from the dHCP using the Unet-preterm model; 4) Apply the same surface extraction and topology correction tool to all the above-mentioned segmented volumes; 5) Visual quality assessment of the obtained segmentations and surfaces. We further describe each processing step below.

**Figure 1:**
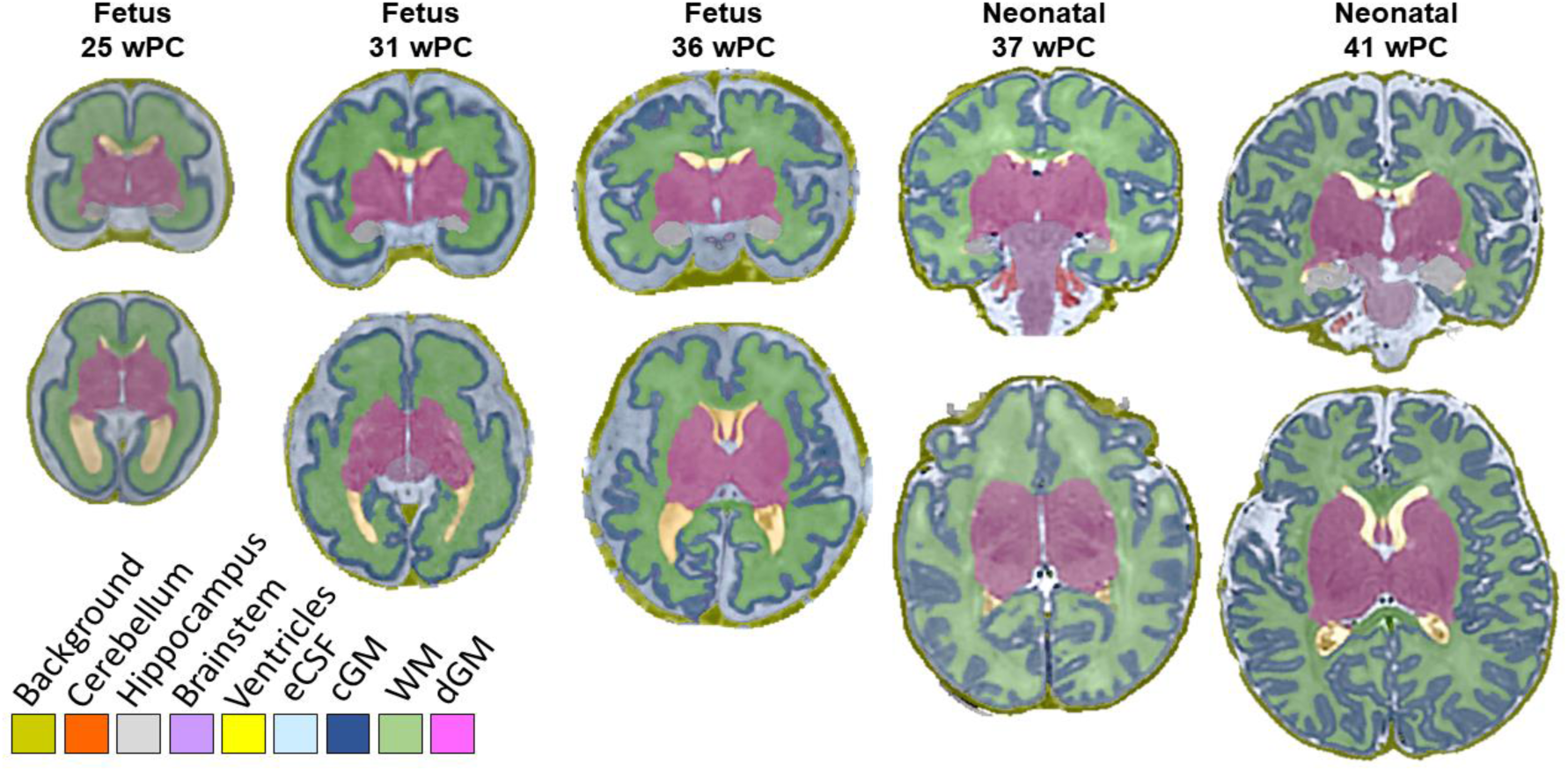
Examples of obtained segmentations for three fetuses and two neonates, illustrating the consistency of the anatomical delineation despite large variations in age and developmental stage. eCSF represents extra-cerebrospinal fluid, cGM represents cortical gray matter, WM represents white matter, and dGM represents deep gray matter.

#### 2.3.1. Training of the segmentation model on the postnatal dataset

The dHCP consortium provides whole brain multi-tissue segmentation maps based on the technique Draw-EM (Makropoulos et al., 2018) that combines a spatial prior and a model of the intensity of the image in order to enforce robustness to variations in image intensity distribution related to brain maturation. This segmentation map includes extra-cerebrospinal fluid (eCSF), cortical gray matter (cGM), white matter (WM), lateral ventricles, deep gray matter (dGM), cerebellum, brainstem, and hippocampus. To avoid the influence of local inaccuracies in the segmentation of the convoluted cortical gray matter resulting from the well known limitations of the atlas-registration approach in Draw-EM, we replaced the initial cortical gray matter label with the cortical ribbon volumetric mask also provided by the dHCP and defined as the voxels located between the inner (white) and the outer (pial) cortical surfaces. We then use the segmentation maps as defined above from the 50 youngest infants of the dHCP to train a first nn-Unet model. The age within the training set ranged from 29.3 to 37.1 weeks post-conception, covering a substantial portion of the fetal period. We denote this model as Unet-preterm in the following sections.

#### 2.3.2. Fine-tuning on the fetal dataset

We used the Unet-preterm model described above to predict the segmentation from the 3D T2w fetal MRI volumes that passed the visual quality assessment. As expected, the accuracy of these segmentations was not always satisfying. We adopted the ‘active learning’ strategy as introduced in Uus et al. (2023) and Budd et al. (2021) by manually refining the initial segmentation predicted by the Unet-preterm model. We specifically screened the highest quality fetal MRI volumes, which is critical for accurate manual delineation of anatomical structures, and selected seven cases showing minimal, unambiguous errors. We obtained optimal quality data and corresponding accurate segmentation for these seven fetuses. We then fine-tuned the Unet-preterm model using these ground truth fetal data following the procedure described in (https://github.com/MIC-DKFZ/nnUNet/blob/master/documentation/pretraining_and_finetuning.md). During this fine-tuning, the model further converges toward an optimal configuration that lies closer to the Unet-preterm model than in the case of training directly from fetal data. This second model is denoted as Unet-fetus. Extensive visual assessment of selected cases for which the prediction from the Unet-preterm was inaccurate confirmed that the fine-tuning on seven fetal data was sufficient to proceed with the next step of our pipeline.

#### 2.3.3. Segmentation of the postnatal dataset using the nn-Unet based model

In order to ensure a consistent image processing across both postnatal and fetal datasets, we also used the Unet-preterm model to predict the segmentation of the remaining subjects from the dHCP dataset. The visual assessment described in section 2.3.5 below confirmed that this segmentation model is more accurate than Draw-EM provided by the dHCP consortium.

#### 2.3.4. Whole brain mesh generation and surface features

In order to extract a surface mesh allowing us to compute surface features, we designed an extended white matter mask from the multi-tissue segmentation by aggregating the labels corresponding to the ventricles, deep gray matter and hippocampus with the white matter. Figure 2 shows examples of cortical surfaces extracted using our pipeline spanning the perinatal period from fetal to postnatal subjects. A 3D global gyration index similar to the one proposed in Batchelor et al. (2002) was computed as the ratio between the white mesh surface area and the area of its convex hull derived directly from the mesh using the SLAM toolbox (<u>brain-slam/slam:Surface anaLysis And Modeling</u>).

**Figure 2:**
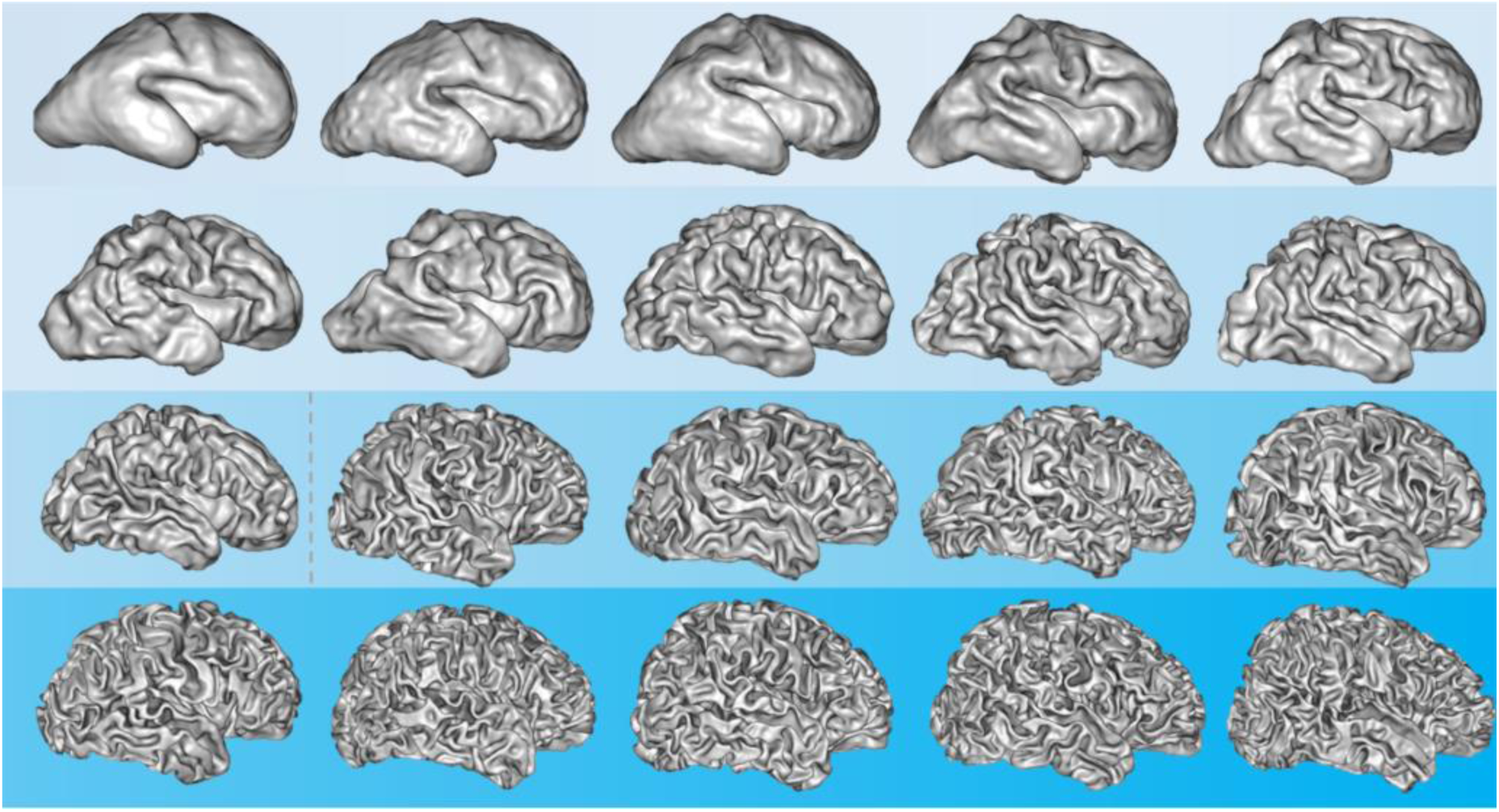
Examples of cortical surfaces obtained using our pipeline for each week post-conception from 26 to 45 weeks (left to right, along each row). Note the increasing folding with age. The dotted gray line represents the transition from fetal to postnatal cortical surfaces.

#### 2.3.5. Quality control (QC)

In order to ensure a high accuracy of the features used in our statistical analyses described below, we conducted an extensive visual assessment of the quality of the images, segmentations and corresponding cortical surfaces. Three raters (G.A, A.M and A.P) scrutinized the 3D reconstructed T2w volumes to identify potential artifacts and anatomical inaccuracies. Regarding T2w postnatal MRI data, we relied on the quality control and the radiological screening performed by experts involved in the dHCP consortium and incorporated postnatal T2w MRI scans that were successfully processed using the full dHCP structural pipeline (Makropoulos et al., 2018). We also controlled the quality of the cortical surface mesh for each participant from both fetal and postnatal datasets. The visual assessment of the cortical surface is efficient for spotting subtle segmentation inaccuracies that would be very hard to detect by inspecting the segmentation in the voxel space.

### 2.4. Statistical analyses

#### 2.4.1. Normative models using the GAMLSS framework

As described in Marquand et al. (2016), the aim of normative modeling is not only to estimate a regression curve representative of an average across a population, but also to estimate centiles of a distribution for each brain phenotype that will serve to quantify the potential deviation from the reference model. In this work, we used the generalized additive model for location, scale and shape (GAMLSS) framework to observe and model the evolution of each brain feature as a function of age as subjects traverse the birth barrier. This is a robust and flexible normative modeling framework introduced in Stasinopoulos and Rigby (2008) that can be fit to non-Gaussian distributions and used to characterize heteroskedasticity and dynamic non-linear growth. We used the implementation provided by Dinga et al. (2021), which is based on the R package presented in Stasinopoulos and Rigby (2008). This implementation of the GAMLSS framework provides automated parameter optimization to obtain optimal fit for inserted data. It is therefore ideal for modeling the median and centiles of non-linear neurodevelopmental trajectories of brain features extracted from neuroimaging data as a function of age (Kyriakopoulou et al., 2017; Bethlehem et al., 2022; Frangou et al., 2022; Bozek et al., 2023). GAMLSS was fit with the Sinh-Arcsinh (SHASH) model distribution for growth as a suitable four parameter distribution corresponding to location, scale, skewness and kurtosis due to its flexibility across different types of data distributions including scenarios of non-normally distributed residuals (Jones and Pewsey, 2009; Dinga et al., 2021).

The GAMLSS model was fit on each of the following whole-brain volumetric features: cortical gray matter, deep gray matter, white matter, cerebellum, brainstem, hippocampi, extra-cerebrospinal fluid (eCSF) and ventricles. Whole-brain gyrification and white matter surface area were also investigated. Quantile intervals were set by the model and computed at [-2σ, ™1σ, median, 1σ, 2σ] in order to better incorporate the data. Model quality of the fit per feature was assessed by computing model diagnostics metrics including worm plots, Q-Q plots and other distribution measures such as skewness, kurtosis and Filliben correlation coefficients. All analyses were done in R using gamlss, nlme and mgcv packages, and plots were generated using ggplots. We used the code provided by Dinga et al. (2021) as a starting point for our statistical analyses (https://github.com/dinga92/gamlss_normative_paper).

#### 2.4.2. Normalized trajectories and velocity

To better visualize and compare the neurodevelopmental trajectories of all features, normalization is required in order to compensate for potentially large variations related to global tissue sizes. In this work, we also normalized the median trajectory of each feature by its maximum, which corresponded to the value recorded at 45 wPC (the oldest age) for all cortical features. These normalized trajectories can thus be collectively interpreted in terms of evolution relative to the highest cortical feature value (achieved at the end of our age range for all features at 45 wPC). In order to further characterize the dynamics of each feature’s cortical trajectory, we further computed their respective first derivatives, which describes the velocity, or growth rate, at each point on the curve.

#### 2.4.3. Feature Proportionality Across Age

To observe inherent maturational dynamics of the cortical features collectively, we calculated the proportion of the intracranial volume (ICV) that each feature represents per week across the perinatal period using the following formula:

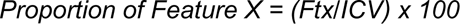

*Ft* represents the cortical feature, *x* represents the type of cortical feature (computed for each of the following = brainstem, cerebellum, cortical gray matter, deep gray matter, white matter, hippocampi, eCSF and ventricles), and *ICV* is the summation of all the volumetric features. This analysis was computed on only volumetric features (i.e. not including surface area and gyrification).

#### 2.4.4. Sensitivity analyses

To assess the robustness of our statistical models we tested for sources of potential influence on the estimated trajectories including the effect of sex and of the type of scanner used for MRI data acquisition.

##### 2.4.4.1. Effect of sex

Recent works such as Studholme et al. (2020) report significant effects of sex on brain growth trajectories even before birth. Including sex as a categorical factor in the model is thus expected, if possible. However, since fetal sex was not systematically recorded in the clinical routine implemented at the hospital of la Timone since 2008, this information is missing in 16% of our fetal dataset. We chose to proceed with a sensitivity analysis instead of excluding this 16% of our fetal population. To do so, we re-ran the GAMLSS models on males and females independently for those participants *with* sex data in order to empirically observe if brain feature continuities were influenced by the effect of sex. GAMLSS models were also re-run including sex as a factor on all subjects with sex data to isolate potential statistical differences in cortical features between males and females.

##### 2.4.4.2. Effect of scanner in fetuses

Another well known confounding factor is the type of scanner used for MRI acquisition, which can affect the image processing algorithm and thus the estimated brain features (Radua et al., 2020). As previously mentioned, we used two separate cohorts to conduct our investigation: the publicly available dHCP dataset for postnatal participants, which only used a 3T scanner, and our local MarsFet dataset for fetal participants, which used both 1.5T and 3T scanners. For each brain feature in the fetal cohort we ran separate linear models for those using a 1.5T scanner and those using a 3T scanner, to empirically observe if their brain feature trajectories were influenced by differences in scanner strength. A linear model was deemed preferable for use on our fetal cohort due to the limited sample size. Linear models were also run on the entire fetal cohort per cortical feature including scanner type as a factor to isolate any statistical differences between fetal subjects scanned with a 1.5T scanner and those scanned with a 3T scanner.

## 3. Results

### 3.1. Data selection and description of final sample

The flow chart recapitulating the data selection process is provided in Figure 3. Regarding the fetal dataset, 806 MRI sessions including a HASTE sequence have been acquired between 2008 and 2021 at the hospital of la Timone. Based on the clinical evaluation from the neuroradiologist, 404 fetal sessions were considered normal. We then applied the exclusion criteria described in section 2.1 and Table 2, which resulted in 300 fetal sessions corresponding to ‘clinical controls with normal MRI’. For each scanning session, we estimated a 3D T2w volume. The visual quality check procedure described in section 2.3.5 resulted in the exclusion of 229 volumes of insufficient quality, leaving 175 3D T2w volumes to be kept in the pipeline. Finally, 25 more cases were excluded based on the visual assessment of the segmentation maps and cortical mesh, resulting in a final sample of 150 MRI sessions (65 males, 61 females, 24 unknown) from 140 healthy fetuses covering an age range between 23.7 and 37 (mean = 31.6; std = 2.5) weeks post-conception.

**Figure 3:**
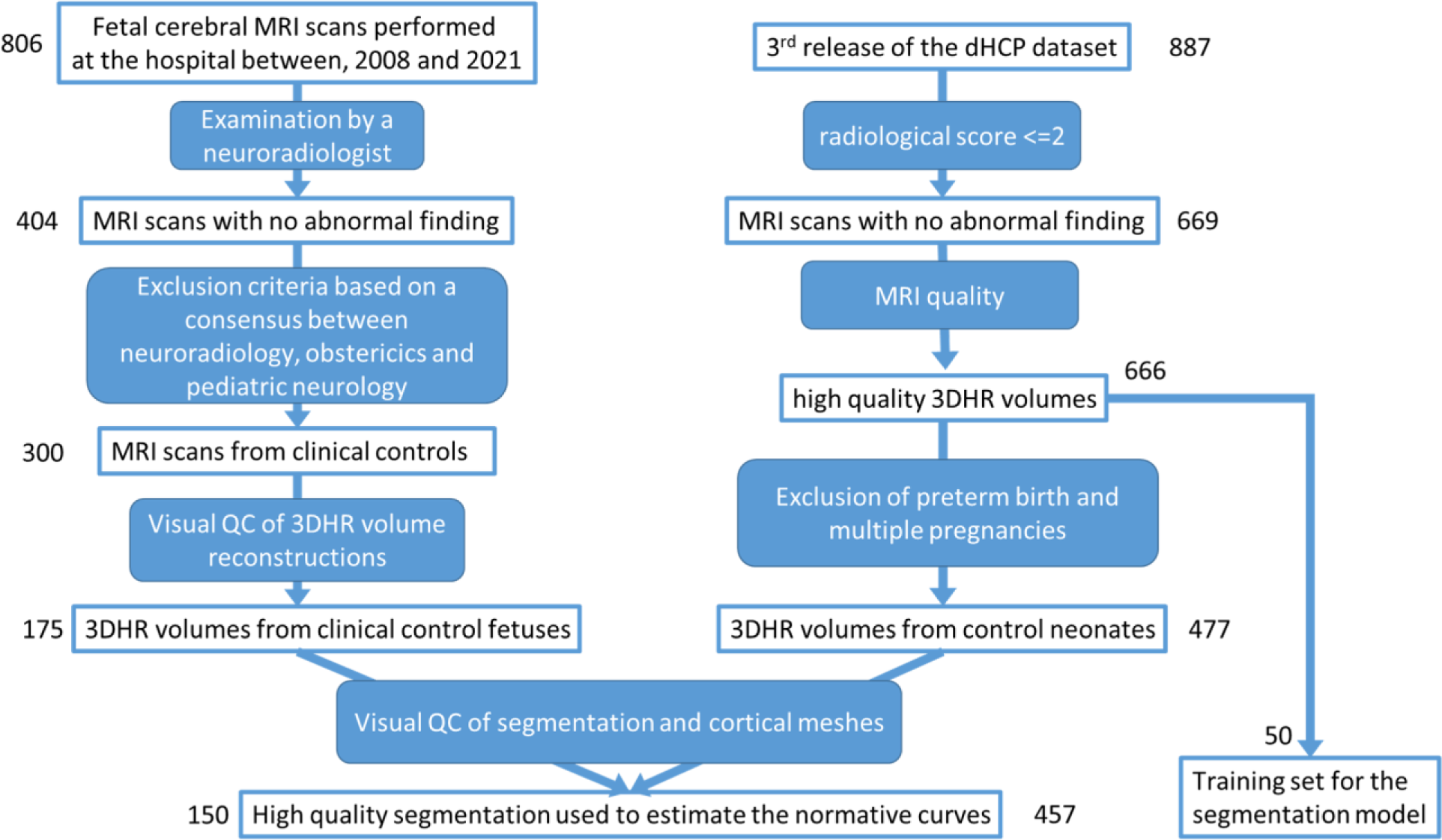
Data selection flowchart for fetal (left) and postnatal (right) sessions.

Regarding the postnatal dataset, 669 out of 887 MRI scanning sessions were selected based on postnatal clinical and imaging exclusion criteria of the dHCP dataset. 666 MRI sessions passed the quality control conducted by the dHCP consortium. After exclusion of multiple pregnancy and preterm birth, we obtained 477 MRI sessions. Lastly, 20 MRI sessions were excluded based on the visual QC of the segmentation and cortical mesh, resulting in a final sample of 457 MRI sessions (248 males, 209 females) from 456 healthy newborn babies covering an age range between 37.4 and 44.9 (mean = 41.5; std = 1.6) weeks post-conception.

Selection from these two cohorts resulted in our final *perinatal* sample which spanned an age range between 23.7 and 44.9 (mean = 39.1; std = 4.7), and included 607 sessions from 596 healthy participants. See Table 3 below for basic demographic information of the perinatal cohort which is composed from the MarsFet and dHCP datasets.

**Table 3:** Basic demographic information for the sessions of the perinatal cohort, which is composed from the MarsFet and dHCP cohorts.

|  | Sex Ratio<br>(M : F : NA) | Scan Age Range<br>(Gestational Weeks) | Scanner Type Ratio<br>(3T : 1.5T) |
| --- | --- | --- | --- |
| Perinatal Cohort | 270 : 313 : 24 | 23.7 – 44.9 (mean = 39.1; std = 4.7) | 541 : 66 |
| Fetal Cohort: *Marsfet* | 65 : 61 : 24 | 23.7 – 37.0 (mean = 31.6; std = 2.5) | 84 : 66 |
| Postnatal Cohort: *dHCP* | 248 : 209 : 0 | 37.4 – 44.9 (mean = 41.5; std = 1.6) | 457 : 0 |

Finally, as described in Section 2.3.1, we selected 50 scanning sessions acquired at earlier ages from the dHCP to train our Unet-preterm segmentation model, which have a scan age range of 29.3 - 37.1 weeks post-conception (34 males, 16 females), and a birth age range of 25.57 - 36.85 weeks post-conception (mean age=32.7, std=3.14).

### 3.2. Normative Modeling Analysis

#### 3.2.1. Model selection

As suggested in Dinga et al. (2021), we ran a 4-model comparison using our data: a simple linear regression model, a GAM homoskedastic model, a GAM heteroskedastic model, and a GAMLSS model. Visual inspection indicated that the GAM heteroskedastic and GAMLSS models were better fit to the data compared to the simpler models (Extended Figure 4-1). This was further validated by computing the Akaike Information Criterion to estimate the quality of fit of each model for each cortical feature (Extended Figure 4-2). GAMLSS outperformed the remaining models in the majority of estimations and was therefore chosen to represent our data in the current investigation.

#### 3.2.2. Normative plots

As expected, all cortical feature trajectories show an increase in volume with age. As illustrated in Figure 4, the median trajectory is almost linear for the total intracranial volume (ICV), brainstem, deep gray matter, and hippocampi volumes. The trajectory follows an exponential pattern for the cerebellum, and, to a lesser extent, the cortical gray matter. A slight inflection is visible for the total white matter volume with a modest decrease in growth following birth. The trajectory for the cortical surface area shows an increase after birth compared to during the fetal period. Remarkably, the development of the aforementioned cortical features follows a smooth median trajectory with maintained variability from early fetal life up until around two months postnatally, without a significant discontinuity at birth.

**Figure 4:**
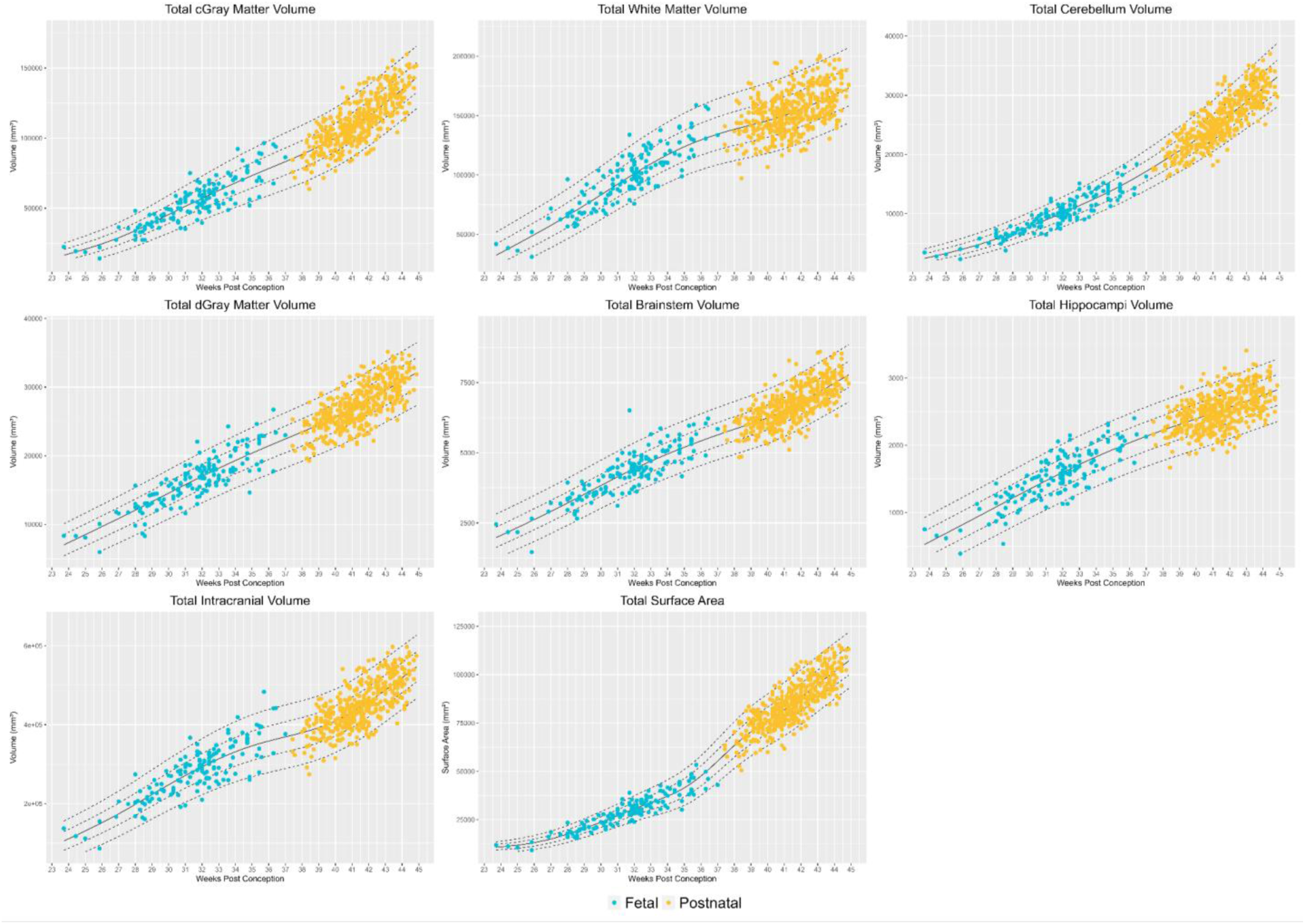
The normative plots of 7 ***continuous*** global volumetric features = Cortical Gray Matter (cGray Matter), White Matter, Cerebellum, Deep Gray Matter (dGray Matter), Brainstem, Hippocampi and intracranial volume; and one surface feature = Total Surface Area. Blue represents the fetal MarsFet cohort and orange represents the postnatal dHCP cohort.

As illustrated in Figure 5A, we observe different patterns for the three remaining measures: the eCSF volume, ventricles volume and gyrification index. These features exhibit a discontinuity either in variance or in value. The strikingly different patterns observed for these measures before and after birth cast doubt on the validity of assuming continuity in estimating perinatal trajectories. To analyze these patterns in more detail, we computed two independent normative models for each of these three measures: one for the fetal period and one for the postnatal period (Figure 5B). Each of these three features showed distinctive patterns. The variability around the median trajectory for ventricles volume is considerably larger during the fetal period than in the postnatal period. Regarding the eCSF volume, the variance is also higher before birth compared to after. As illustrated on the ‘two models’ plot (Figure 5B), the eCSF volume variance shows a constant increase with age for the fetuses, while the postnatal variance is much less. The comparison between the median eCSF trajectory from a single (Figure 5A) versus a ‘two models’ (Figure 5B) experiment suggests that the transition might still be continuous, with a plateau around birth. Lastly, the trajectory obtained for the global gyrification index is striking, with a relatively maintained variability but a sharp transition from fetal to postnatal life. In addition to this acceleration, the slope of the median curve is higher after birth than before.

**Figure 5:**
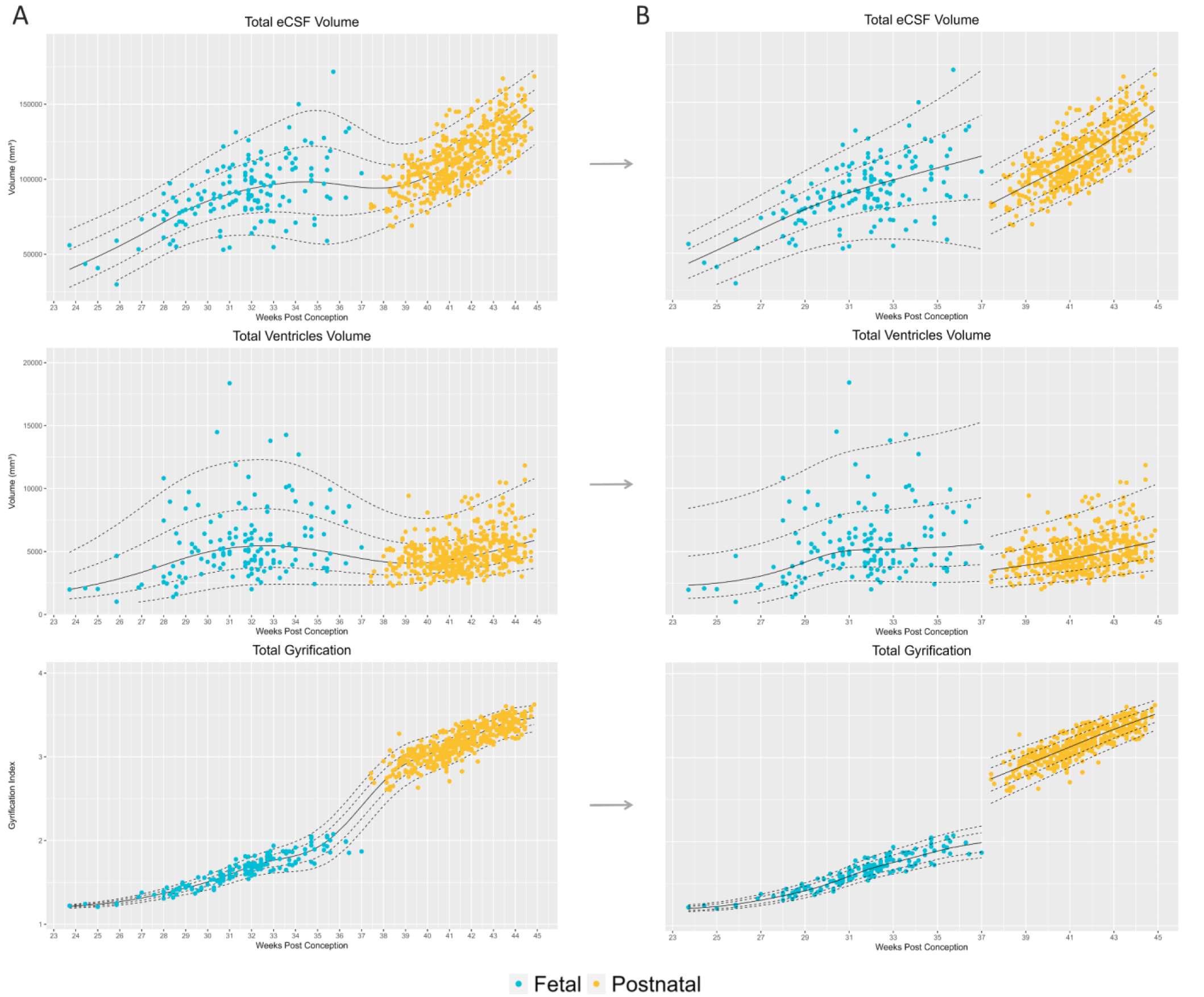
The normative plots of 2 volumetric features = Total eCSF and Ventricles, which exhibit **discontinuity** in *variability* before and after birth; and one surface feature = Total Gyrification, which exhibits **discontinuity** in *growth* before and after birth. In panel A one GAMLSS model was computed for the entire perinatal population, and in panel B two separate GAMLSS plots were computed per participant type (i.e, fetal and postnatal). This was done in order to comprehensively visualize inherent discontinuities. Blue represents the fetal MarsFet cohort and orange represents the postnatal dHCP cohort.

We followed the procedures proposed in Dinga et al. (2021) and Bethlehem et al. (2022) to evaluate model fit by computing Q-Q plots, worm plots and other distribution metrics such as skewness, kurtosis and Filliben correlation coefficients, per feature. Detailed model diagnostics are provided in Extended Figures 4-3, 4-4 and 4-5.

#### 3.2.3. Normalized trajectories and derivatives

To conduct a proper visual comparison across the normalized trajectories exhibited in Figures 4 and 5, all cortical feature median trajectories can be superimposed onto a single plot. As described in section 2.4.2, we used the maximal value of each trajectory for normalization, which constrains all trajectories to reach a maximum value of 1 at 45 wPC. By comparing the normalized trajectories across features, the smoothness for all features except eCSF, ventricles and gyrification is clear. Normalization also enables the comparison of the total growth of continuous features over this developmental period quantified by the difference between the lowest value and 1 (highest value) in each cortical feature. We observe that the total cerebellum volume exhibits the largest total growth with a value of 92.6%, while the total brainstem volume shows the lowest total growth with a value of 74.6% (Figure 6, top). The plots illustrating velocity, or growth rate, confirm more complex trajectories for the eCSF, ventricles and gyrification, with higher variations along their respective trajectories compared to the other features (Figure 6, bottom).

**Figure 6:**
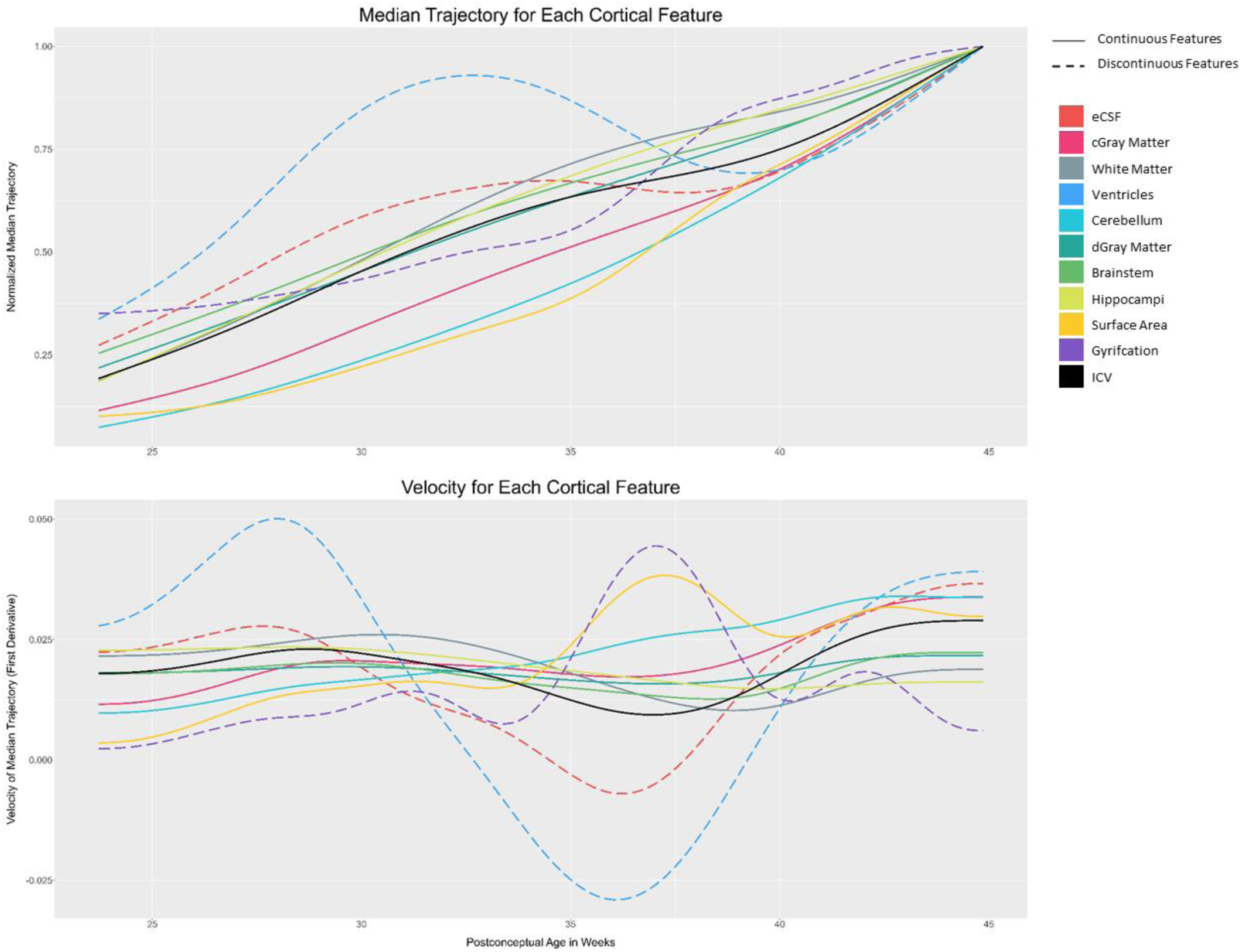
Normalized trajectories (top) and their respective velocities, or growth rates, (bottom) for each cortical feature.

Of note, the median trajectory and velocity for the total surface area show an intermediate level of complexity. In terms of complexity and linearity of the median trajectory, the trajectory pattern for this feature lies between the continuous and discontinuous features (i.e., slightly less linear compared to tissues showing a clearly continuous trajectory, but much more smooth compared to the eCSF, ventricles and gyrification index). Regarding the velocity, the surface area shows a clear burst in growth rate, similar to the pattern elicited by the gyrification. Referring back to Figure 4, we conclude that the trajectory for the total surface area does show a subtle inflexion around birth, but not a discontinuity as observed for eCSF, ventricles and gyrification index.

#### 3.2.4. Feature proportions across age

To investigate how cortical features change within the ICV during development, we plotted the proportion (of the ICV) of each volumetric feature per week across the perinatal period. As shown on Figure 7, we found that most features maintained their proportions across this period with the eCSF showing the most drastic and sharp change in proportion across the perinatal period. Specifically, the eCSF encompassed a larger proportion of the ICV in the fetal stage, with a drastic drop at birth and a smaller proportion in the postnatal period.

**Figure 7:**
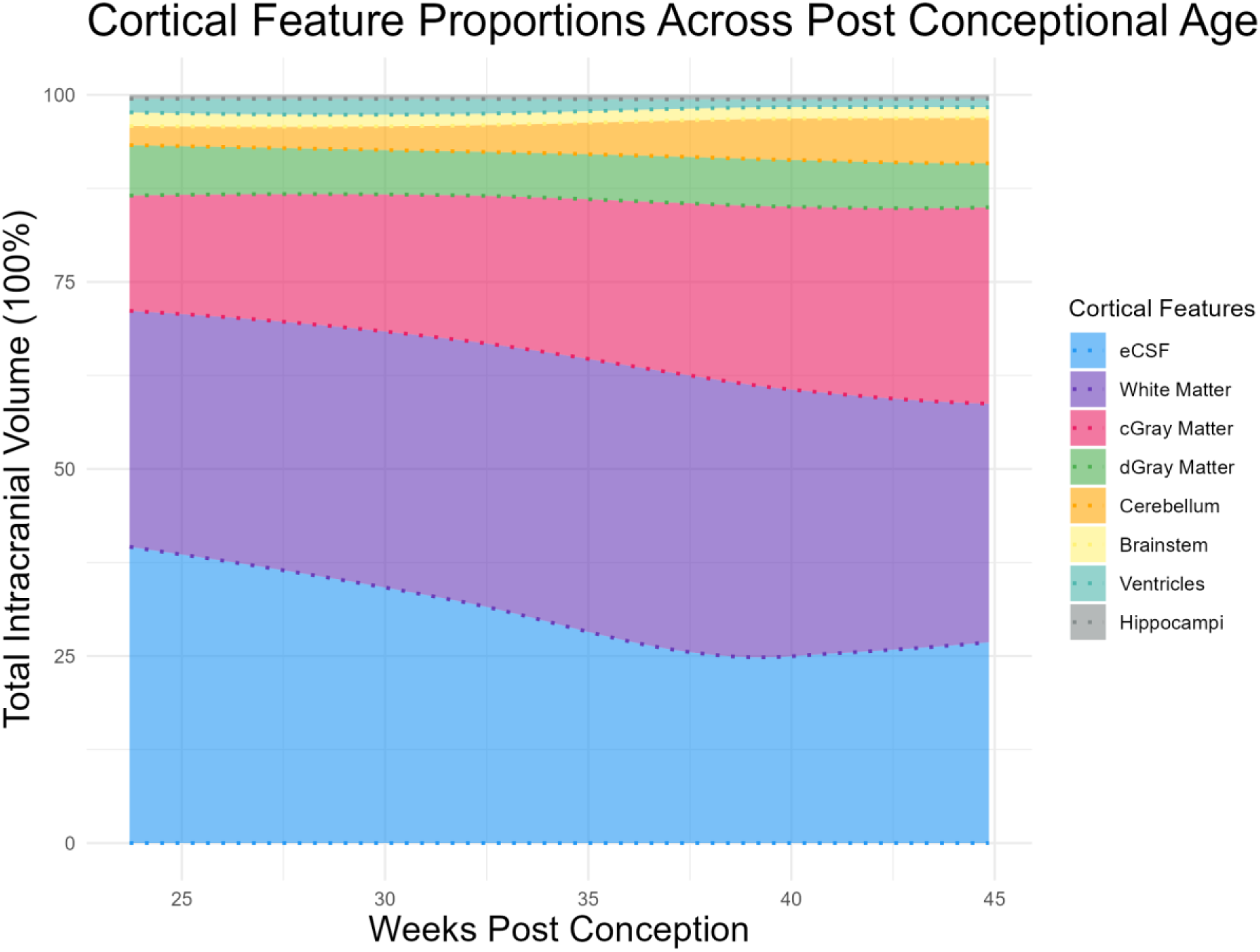
Changes in cortical feature proportions were observed across post-conceptional age with marked variations in the eCSF. Each layer represents one cortical feature. The y-axis represents the percentage of the ICV that each volumetric feature encompasses per week (summing up to 100% at each week).

### 3.3. Sex effects

For the main analysis focusing on the transition from fetal to postnatal life, we did not stratify our cohort by sex due to incomplete sex data. However, to better understand the role of sex in these normative models, we selectively analyzed participants with sex data (n = 583, 313 males and 270 females) to empirically determine potential sex-specific variations. We conducted a GAMLSS model for each feature, including sex as a covariate to isolate statistical differences. We found highly statistically significant differences in sex in all the volumetric features and in the surface area. All values were Bonferroni corrected. These statistical differences are likely due to established variances in intracranial brain volume between males and females. Upon adjusting for intracranial volume, sex differences were observed for white matter (p = 0.007), cerebellum (p = 8.80 × 10⁷), deep gray matter (p = 0.0057) and brainstem (p = 8.16 × 10⁹) volumes (Figure 8; Extended Figure 8-1).

**Figure 8:**
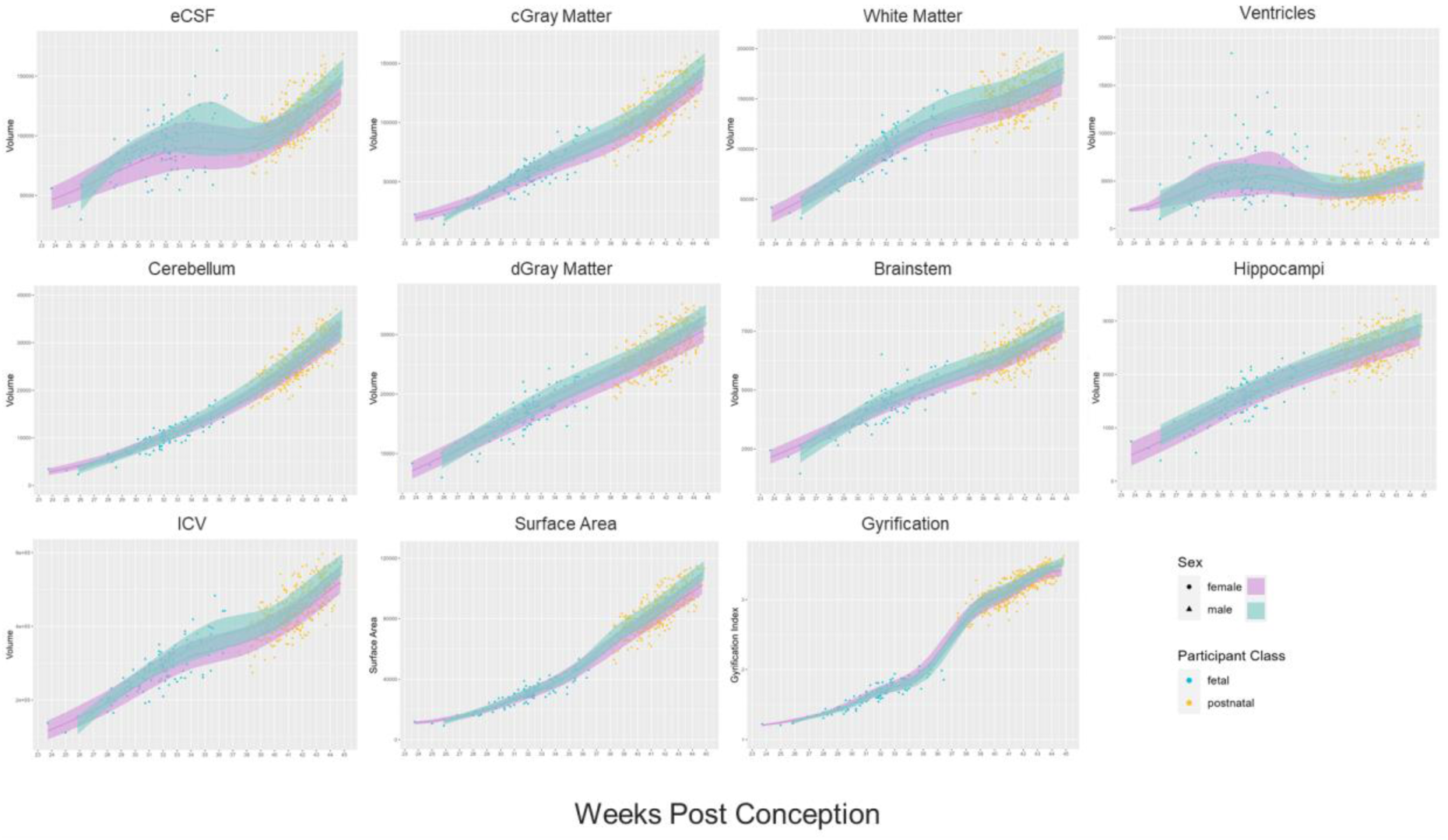
Overlapping normative plots for males and females per feature illustrating the globally consistent normative pattern that can be seen in both sexes. This confirms that any significant sex differences do not affect the continuity and pattern of these neurodevelopmental trajectories.

While these effects related to sex deserve further investigation, they do not compromise our primary goal of investigating the continuity of features using normative modeling across the entire cohort, which included both males and females. We observed similar developmental trajectories between males and females, as shown in Figure 8. More specifically, for both females and males, we observed a clear continuity for all tissues except for eCSF, ventricles and gyrification index as observed in the entire sample (Figures 4 and 5).

### 3.4. Scanner effects

Only one scanner was used for the postnatal dHCP cohort. The MarsFet cohort, however, used two types of scanners, a 3T Skyra and a 1.5T SymphonyTim. We assessed the potential scanner effects in the fetal cohort using simpler linear models since the sample size was too low for proper estimation using a full GAMLSS model. We found significant differences in volume between the two scanners for fetuses in the cortical gray matter (p = 2.16 × 10¹⁶), brainstem (p = 0.031) and the hippocampi (p = 4.40 × 10⁷) (Table 4; Extended Table 4-1). All values were Bonferroni corrected. These effects might be attributed to residual influence of the acquisition sequence and settings on the estimated volumes.

**Table 4:** A summary of the statistical differences in each feature between the 1.5T and 3T scanners in the fetal population. All values are Bonferroni corrected. Due to the limited sample size a linear model was deemed sufficient to compare scanner effects in the fetal population. Values marked with a ‘*’ are significant.

| Cortical Feature | Scanner Effects (p-values) |
| --- | --- |
| eCSF | 0.0705 |
| cGray Matter | $< 2e-16^*$ |
| White Matter | 10.65 |
| Ventricles | 0.547 |
| Cerebellum | 0.182 |
| dGray Matter | 0.0693 |
| Brainstem | $0.0314^*$ |
| Hippocampi | $4.40e-07^*$ |
| White Surface Area | 7.755 |
| Gyrification | 0.135 |
| ICV | 0.092 |

## 4. Discussion

Transitioning from an intra- to an extrauterine environment requires a significant adaptation affecting the entire fetal/newborn body, including the brain. To the best of our knowledge, this is the first neurodevelopmental study targeting the cortical trajectory of perinatal subjects crossing the birth barrier, including MRIs from both fetal and postnatal subjects, with homogenized preprocessing techniques. Furthermore, for global surface area and for several volumetric features, including total cortical gray matter, white matter, brainstem, cerebellum and hippocampi, trajectories follow a continuity as participants move from a fetal to a postnatal environment. Three features, however, exhibit a strikingly abrupt and discontinuous shift as they cross the birth barrier: the extra-cerebrospinal fluid (eCSF), ventricles and gyrification. This study stands as the first to establish that not all cortical features undergo the same developmental change at birth.

### 4.1. Position with respect to previously published normative curves

The largest-scale study thus far, and the only one that we are aware of combining fetal and postnatal datasets, is Bethlehem et al. (2022). They used a cohort of over 100 000 subjects to track trajectories of cerebral volumes and surface area throughout life starting at 115 days post-conception up to 100 years of age using a GAMLSS model. They demonstrated how total gray matter volume, white matter volume, subcortical volume and surface area rapidly increase from the fetal stage until childhood followed by a slow and steady decline until the end of life, albeit with different maxima and trajectory patterns. Though an impressive study, an important limitation is their lack of unified image processing tools, especially for the perinatal period. They reported disruptions in most cortical features as participants traverse from an in-utero to ex-utero environment. Homogenizing our methods across subjects addresses these limitations (Extended Figure 4-6) and better characterizes perinatal neurodevelopment since we report no discontinuities in the perinatal period in the aforementioned features (Figure 4). In particular, we show that surface area follows a continuous trajectory, contrary to what was shown in Bethlehem et al. (2022). Surface area is a particularly critical feature in assessing neurodevelopmental normality, thus properly characterizing it is essential in the identification of healthy participants and biomarkers linked to neurodevelopmental disorders (Kline et al., 2020; Guardiola-Ripoll et al., 2023).

Many other studies have investigated cortical normative neurodevelopment, but often focused on either fetal or postnatal subjects separately. Those studying the fetal population reported increases across age in surface and volumetric features, small variabilities in surface area, cortical, subcortical and cerebellar volumes, and large variabilities for the eCSF and ventricles (Kyriakopoulou et al., 2017; Studholme et al., 2020; Story et al., 2021), all of which we described in our study. Furthermore, they reported an exponential pattern in cerebellar neurodevelopment, and a rapid increase followed by a steady decline in eCSF volume at the end of the fetal period, again identical to our reported patterns. Postnatal studies covering the same age range as our postnatal population (37 to 45 weeks post-conception, born at-term and healthy) are much less common in the literature. One comparable study is that of Huang et al. (2022), claiming to map perinatal cortical surface area from 29 weeks post-conception to 2 years of age. They however included premature subjects to represent ages below 37 weeks post-conception, which introduces important limitations (Chau et al., 2013; Dimitrova et al., 2021). Nonetheless, in the immediate postnatal period, they describe a linear relationship between post-conceptional age and surface area, similar to what is reported in the literature and to what we observed in our results (Jha et al., 2019; Alex et al., 2023).

### 4.2. Discontinuity of Gyrification, eCSF and Ventricles Volumes

We observed three features exhibiting a sharp transition at birth: the total eCSF, ventricles and gyrification. The ventricles and eCSF are filled with CSF which partakes in a plethora of roles including mechanical and immunological protection, homeostasis, delivery of neural growth signaling molecules, neurotoxic waste elimination and regulation of brain growth via positive hydrostatic pressure (Desmond and Jacobson, 1977; Purves et al., 2001; Johanson et al., 2008; Kapoor et al., 2008; Iliff et al., 2012; Xie et al., 2013). CSF constituents vary according to age, from the embryonic phase up until adulthood, and experience an abrupt drop in protein concentration after birth (Shah et al., 2011; Gato et al., 2014). For both eCSF and ventricle volumes, we report high variance during the fetal phase followed by a notable drop in variability in the postnatal phase, reflecting many investigations (Andescavage et al., 2016b; Kyriakopoulou et al., 2017; Story et al., 2021). We also showed that relative eCSF volumes largely decrease after birth, which is a widely published radiological observation (Gholipour et al., 2017; Li et al., 2019; Dubois et al., 2021; Nagaraj and Kline-Fath, 2022)(Figures 1 and 7). This is corroborated in Lefèvre et al. (2016) where the ratios of eCSF and ventricle volumes are higher in fetuses compared to premature postnatal subjects of the same age. This enforces the interpretation that the event of birth itself causes this discontinuity and not the age.

The third feature reporting discontinuity is global gyrification. Gyrification is known to be a genetically determined feature that begins forming early in the gestational period (10 – 15 weeks post-conception)(Chi et al., 1977; Zilles et al., 2013). An essential role of the gyrification is to draw regions of connectivity closer to one another to decrease action potential transit time and in turn increase overall brain function efficiency (Essen, 1997; White et al., 2010; Gautam et al., 2015). The jump in gyrification that we report after birth (Figures 1 and 5) is consistent with radiological observations (Gholipour et al., 2017; Li et al., 2019; Dubois et al., 2021), and could be linked to the crucial increase in brain stimulation after birth attributed to abrupt and intense sensory stimulation from exposure to the extrauterine environment (Polese et al., 2022). Furthermore, we confirm the preliminary results of Lefèvre et al. (2016) that reported more pronounced folding in premature infants compared to fetuses at the same age. This again enforces the idea that the event of birth itself and not the age is linked to this discontinuity.

A multifaceted array of neuro-ontogenetic processes beginning shortly after conception are responsible for the maturing brain including neurulation, neurogenesis, synaptogenesis, pruning, myelination and neuronal migration (Giedd, 1999; Tau and Peterson, 2010; Kostović et al., 2019). Neuronal migration is the process in which neurons travel along the cortex to reach their final destination in the brain, reportedly guided by CSF flow, and typically halts and/or slows down at birth (Purves et al., 2001; Sawamoto et al., 2006). Disruption of this process leads to *neuronal migration disorders* that are largely based on folding malformations (Copp and Harding, 1999). Additionally, the completion of neuronal migration has been linked to matured gyrification, as corroborated by human and animal models (Borrell and Götz, 2014; Tallinen et al., 2016; Kroenke and Bayly, 2018). Since we report noticeable changes in gyrification, ventricles and eCSF at birth, and that likewise neural migration is linked to these cortical features and is a process that itself is affected by birth, further study is warranted into this complex relationship.

The multi-functionality of the eCSF and ventricles, coupled with the genetic impact on gyrification, underscores the importance of their early quantitative analysis. These traits are associated since the CSF is linked to the physical and cellular evolution of the cortex, including and leading up to the eventual development of cortical folds (Magnotta et al., 1999; White et al., 2010; Gato et al., 2014). The fact that these cortical traits are so impacted when crossing the birth barrier indicates possible physiological and/or mechanical factors linked to the womb, the maternal or the sudden external environment.

### 4.3. Limitations

One limitation in this study is the smaller size of the fetal dataset compared to the postnatal cohort. Nonetheless, the GAMLSS model performs robustly enough to account for this, and to properly capture inherent patterns. Furthermore, we would need to map pathological cases onto our trajectories to obtain a better sense of both typical and atypical neurodevelopment. Also, subjects were labeled as typically developing through the inclusion of normal looking scans, absence of maternal medication, no history of illness, and no reported perinatal complications, however a more accurate categorization could be obtained using neurobehavioral assessments starting at 3 years old. Finally, the cross-sectional nature of our current datasets limits the interpretation of brain maturation dynamics to the population and not the individual level.

### 4.4. Conclusions

In conclusion, by unifying data from fetal and postnatal subjects, our study sheds light on the continuous neurodevelopmental trajectory during the perinatal period, addressing a crucial gap in existing knowledge. We show that cortical features can follow either a continuous or discontinuous pattern within the perinatal period as fetal subjects traverse the birth barrier to become postnatal subjects. The drastic pattern change around birth of certain cortical features could have physiological or mechanical influences from either the intrauterine or extrauterine environment. These original observations warrant further study to better understand trajectory dynamics at the physiological and neurobehavioral level.

## Acknowledgments

The research leading to these results has been supported by the ANR SulcalGRIDS Project (Grant ANR-19-CE45-0014) and the ERA-NET NEURON MULTI-FACT Project (Grant ANR-21-NEU2-0005), funded by the French National Research Agency. Centre de Calcul Intensif d’Aix-Marseille is acknowledged for granting access to its high performance computing resources.

