## Supplemental Data for "Normative models combining fetal and postnatal MRI data to characterize neurodevelopmental trajectories during the transition from in- to ex-utero"

### Extended Data

Extended Data Format → Extended Figure X-i; Figure X refers to a figure in the main text, -i refers to each supplemental element that corresponds to this figure. I.e, Extended Figure 4-2 refers to the second supplementary element (could be a figure or table) that supports Figure 4 from the main text. Extended figures/tables are listed in chronological order.

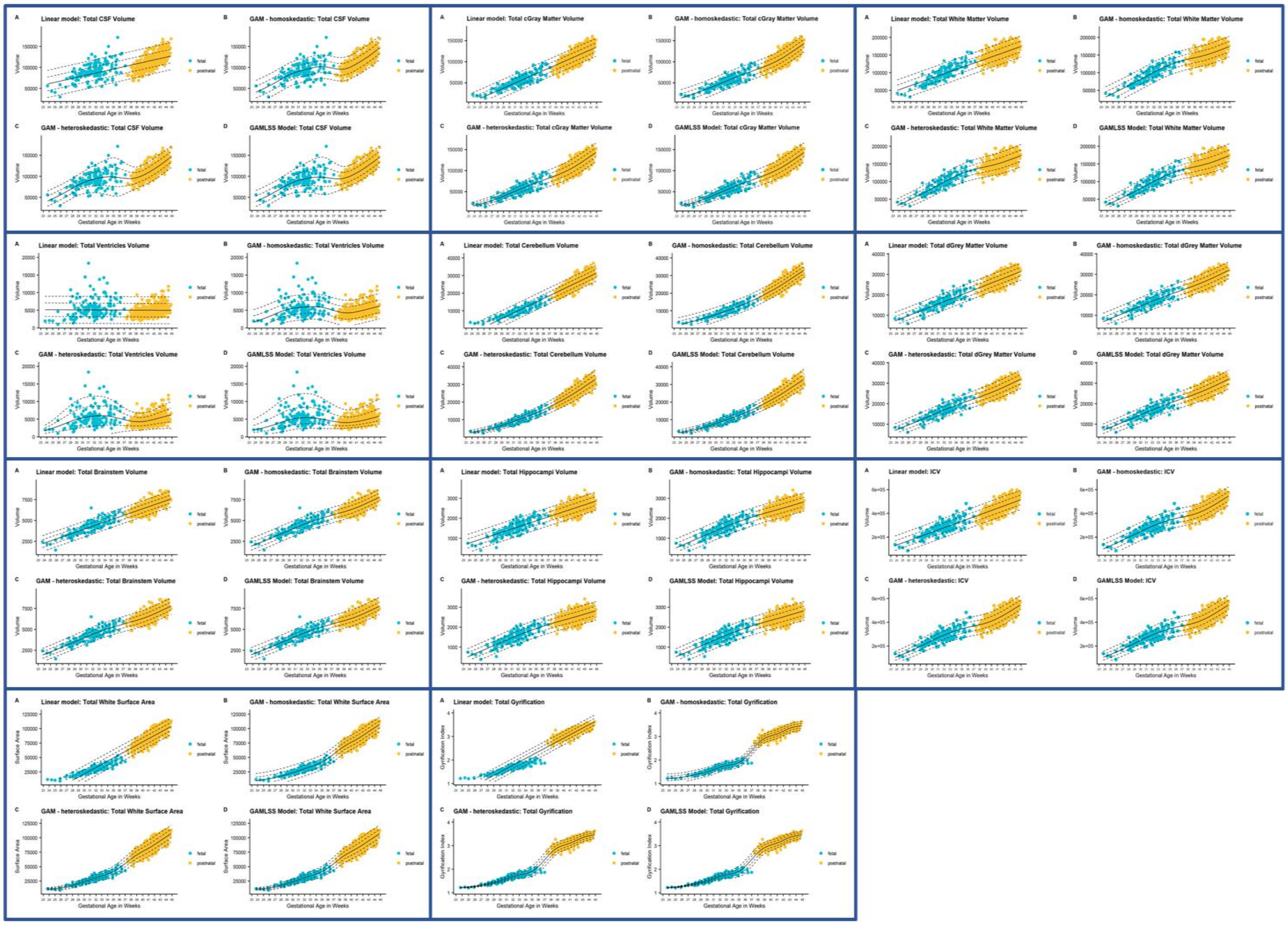


**Extended Figure 4-1:** A 4-model evaluation per cortical feature that tests which model best fits the data.

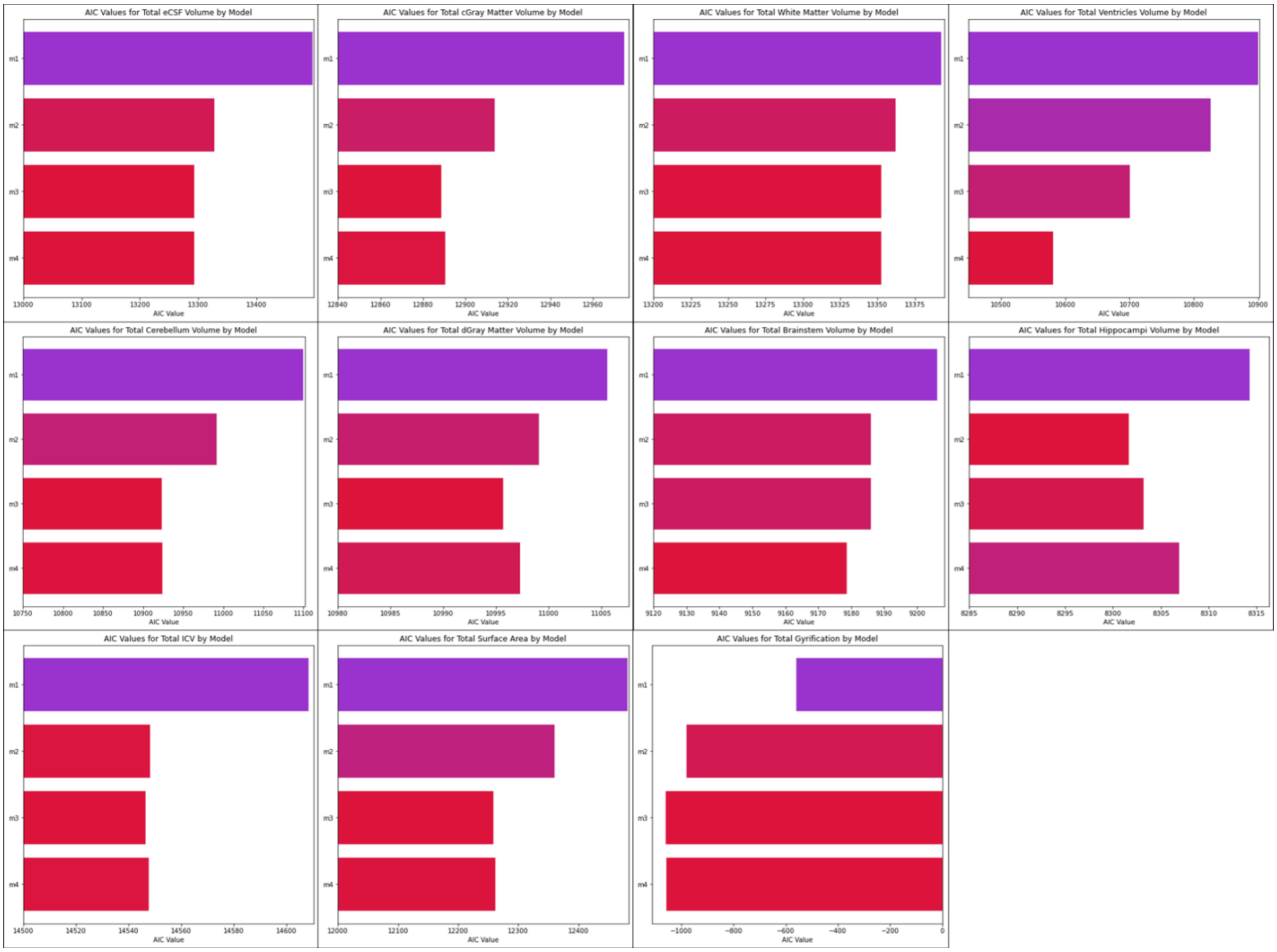


**Extended Figure 4-2:** Akaike Information Criterion plots per cortical feature. M1 through M4 represent the four regression models tested on the data: M1 - a simple linear regression model, M2 - a GAM homoskedastic model, M3 - a GAM heteroskedastic model, and M4 - a GAMLSS model. The GAM heteroskedastic and GAMLSS models generally best fit our data.

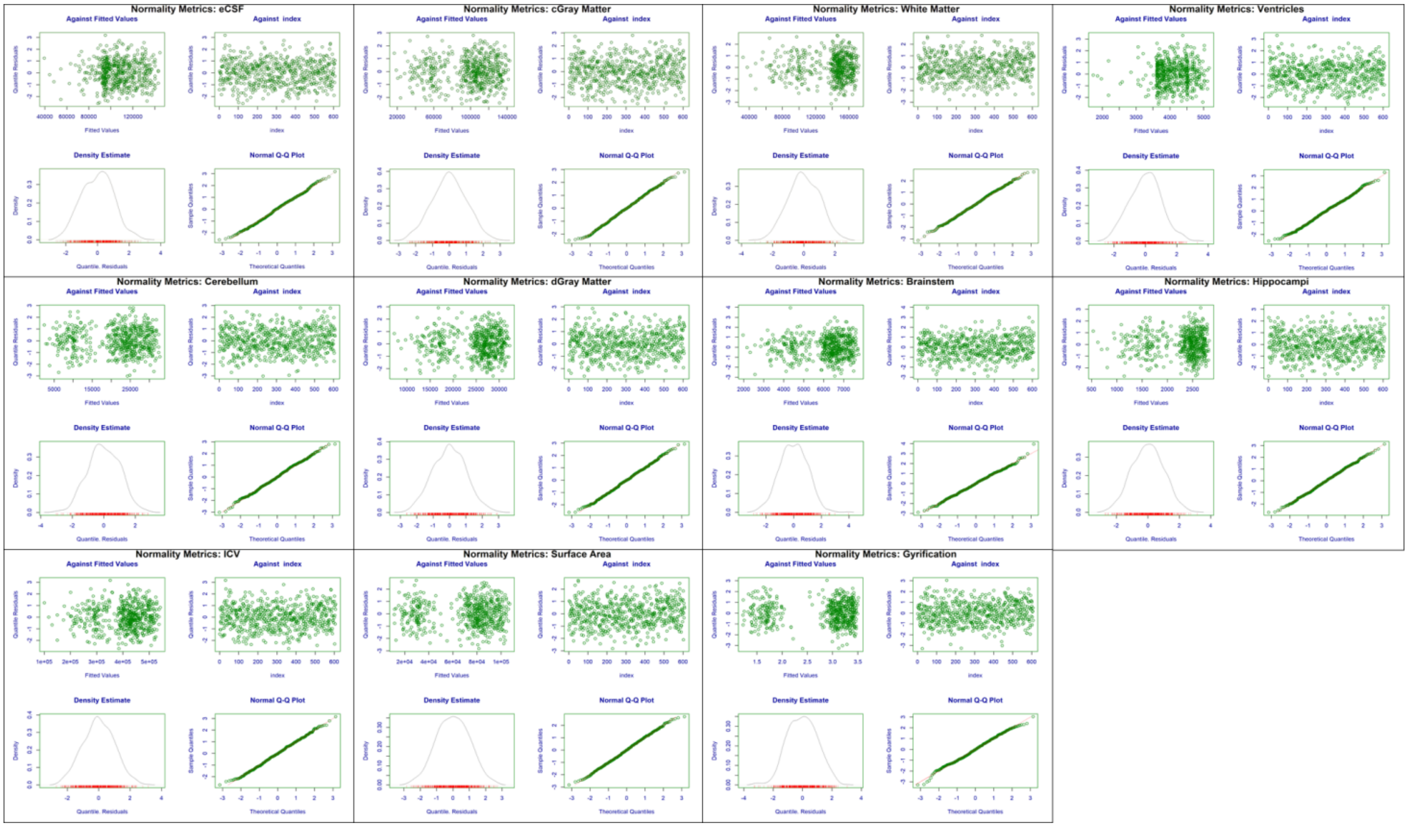


**Extended Figure 4-3:** Model diagnostic plots summarizing normally fitted residuals for each cortical feature, including traditional Q-Q plots.

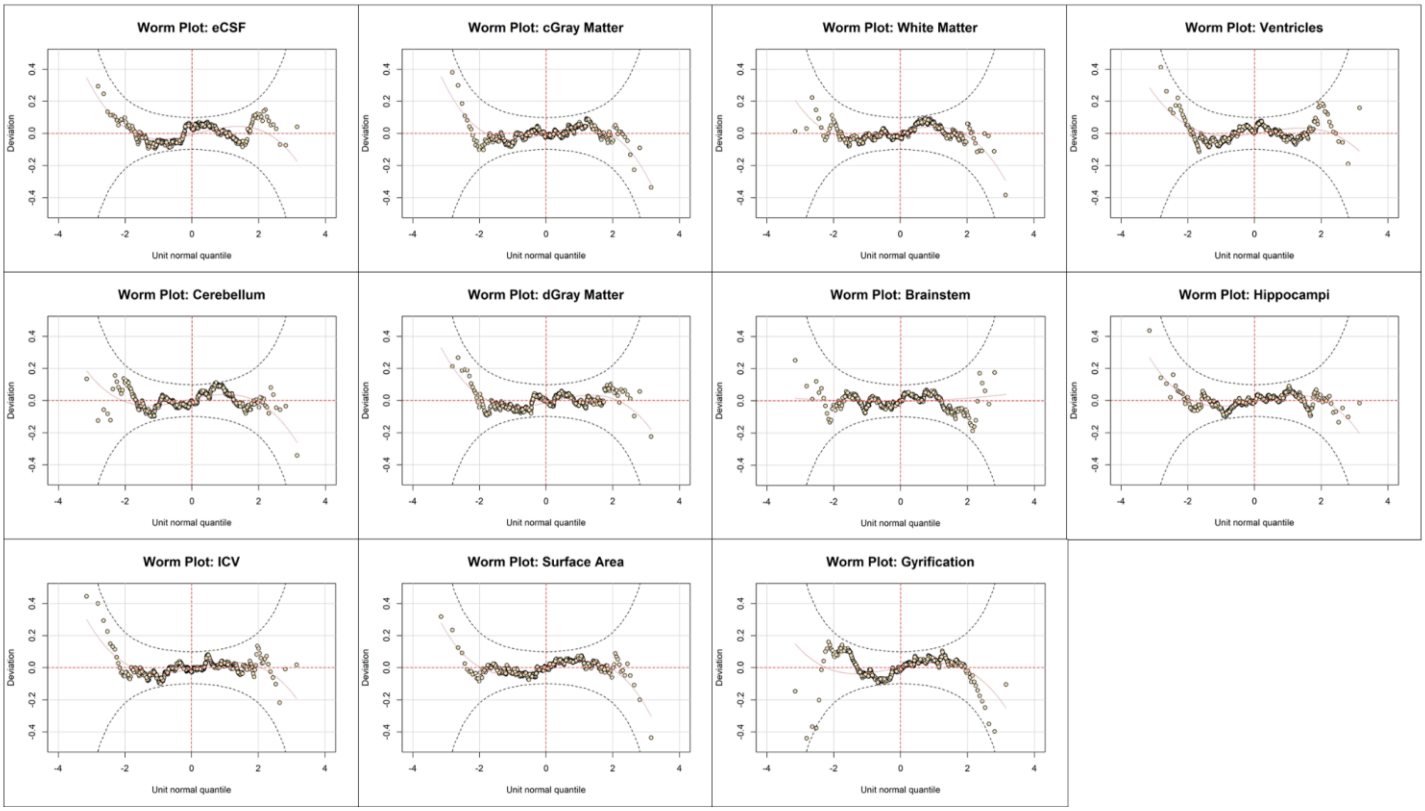


**Extended Figure 4-4:** Further visualization of normality in the form of worm plots for each cortical feature.

| Cortical Feature | Skewness Coefficient | Kurtosis Coefficient | Filliben Correlation Coefficient | Pseudo $R^2$ |
|---|---|---|---|---|
| eCSF | 0.0466761 | 2.715548 | 0.9983508 | 0.6190483 |
| cGray Matter | 0.004863061 | 2.678101 | 0.9987367 | 0.8932966 |
| White Matter | -0.02868258 | 2.705549 | 0.9990697 | 0.7831415 |
| Ventricles | 0.04364758 | 2.811958 | 0.9984278 | 0.2184798 |
| Cerebellum | -0.02714716 | 2.729058 | 0.9987511 | 0.9356335 |
| dGray Matter | 0.0429448 | 2.729334 | 0.9988421 | 0.845485 |
| Brainstem | 0.03024006 | 3.037382 | 0.998564 | 0.8472051 |
| Hippocampi | 0.01777209 | 2.723238 | 0.9991907 | 0.7689623 |
| Surface Area | -0.007055903 | 2.676899 | 0.9991229 | 0.9475578 |
| Gyrification | -0.03652641 | 2.796868 | 0.9978822 | 0.9591794 |
| ICV | 0.02805652 | 2.735637 | 0.9989635 | 0.8323129 |

**Extended Figure 4-5:** A table displaying residual distribution metrics per cortical feature as well as a pseudo $R^2$ value, which is suitable for the GAMLSS regression.

#### Sex Effects

| Cortical Feature | Sex Effect (*p-value*) | Volume Adjusted Sex Effect (*p-value*) |
|---|---|---|
| eCSF | 6.03e-11* | 1.30 |
| cGray Matter | <2e-16* | 4.61 |
| White Matter | <2e-16* | 0.007* |
| Ventricles | 9.66e-04* | 0.71 |
| Cerebellum | 1.09e-03* | 8.80E-07* |
| dGray Matter | <2e-16* | 0.0057* |
| Brainstem | 4.73e-11* | 8.16E-09* |
| Hippocampi | 2.29e-14* | 0.45 |
| White Surface Area | 7.42e-08* | 7.42e-08*† |
| Gyrification | 0.329 | 0.329† |
| ICV | <2e-16 | <2e-16† |

**Extended Figure 8-1:** A table displaying the statistical differences between sex per cortical feature value. Significant differences in sex per volume-adjusted feature and effect sizes are also reported. All values are Bonferroni corrected. Values marked with a '*' are significant. Values marked with a '†' cannot be volume adjusted.

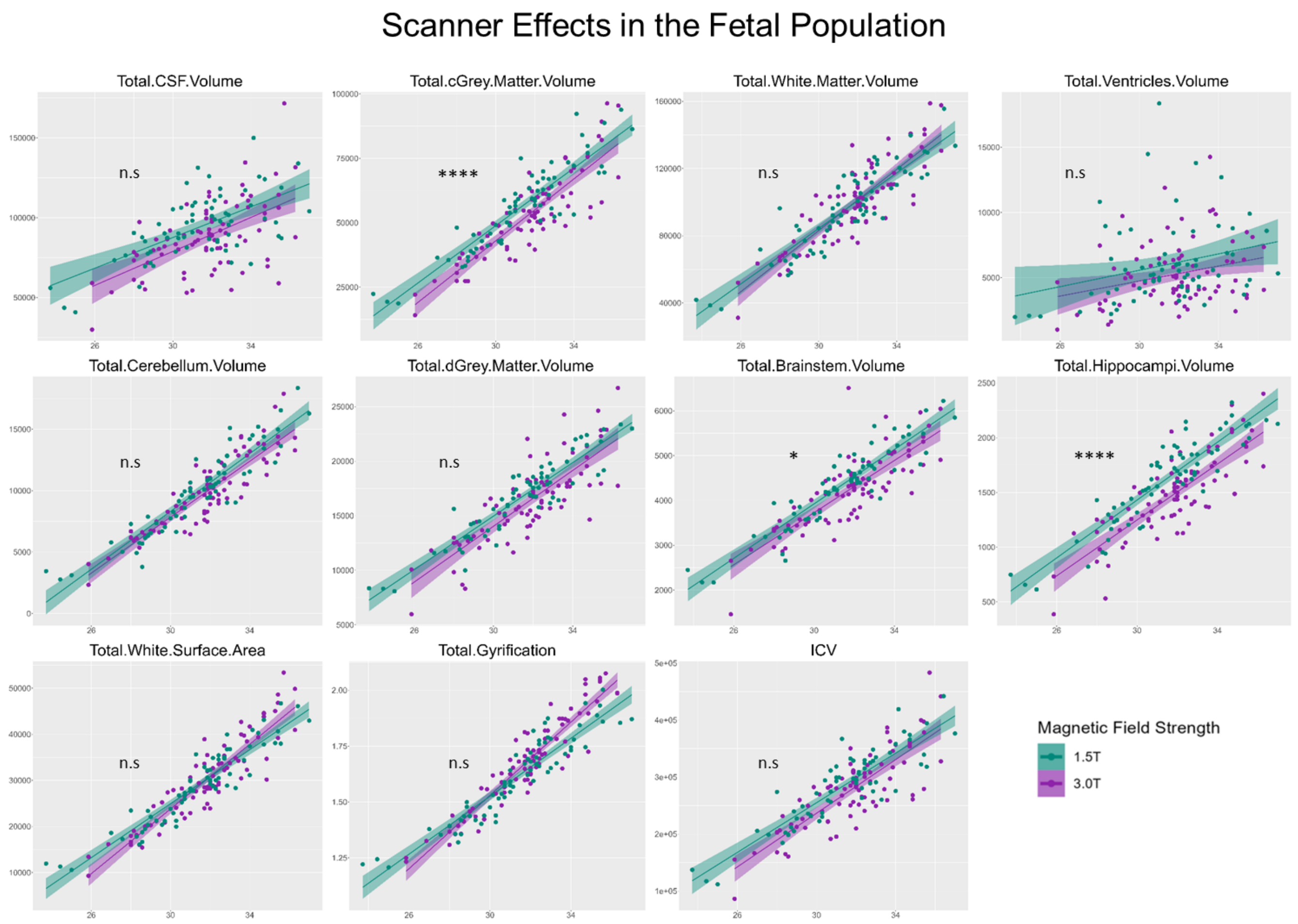


**Extended Table 4-1:** Linear models illustrating the trajectories of fetal subjects that were scanned with a 1.5T and those scanned with a 3T scanner in each cortical feature across post-conceptional age. Cortical features with a significant difference between scanner types are indicated with a '*'. Those not significant are denoted with 'n.s'.

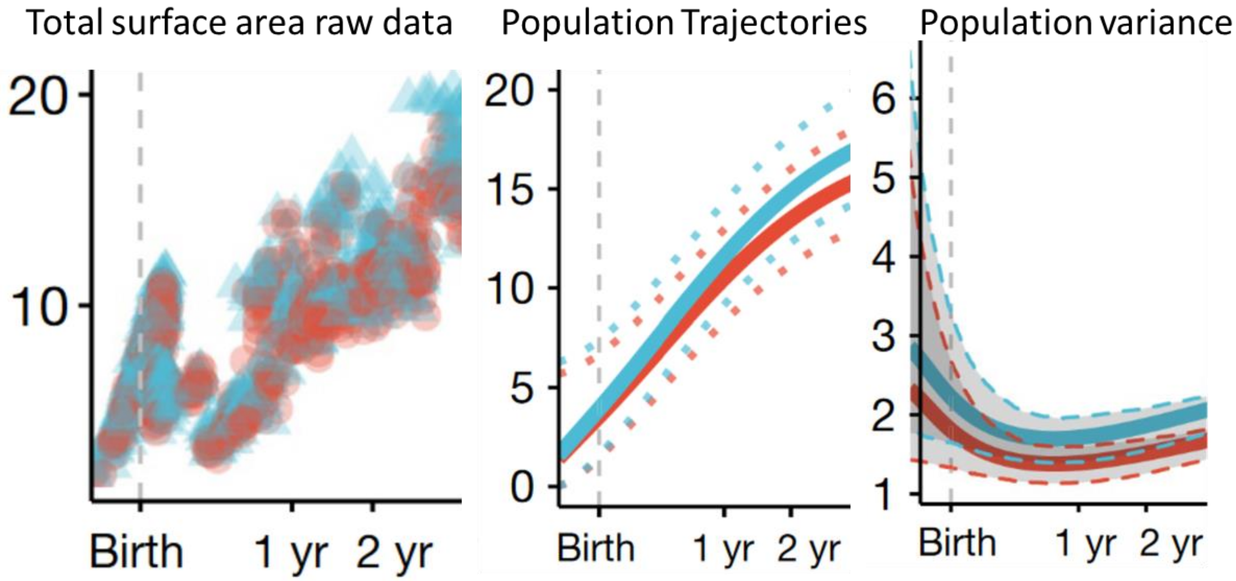


**Extended Figure 4-6:** A zoomed portion of Figure 2 in Bethlehem et al., 2020, which is distributed under the terms of the Creative Commons CC BY license (https://creativecommons.org/licenses/by/4.0/). We illustrate here the lack of uniformization of the image processing tools in the perinatal period from Bethlehem et al. (2020)s, using the measure of total surface area as an example. The lack of uniformization in image processing tools across cohorts explains the large proportion of the variations observed in the raw data (left), the large centile ranges (middle) and high population variance (right) reported on the perinatal period.
